# Heterosis of Fitness and Phenotypic Variance in the Evolution of Diploid Gene Regulatory Network

**DOI:** 10.1101/2021.11.21.469482

**Authors:** Kenji Okubo, Kunihiko Kaneko

**Affiliations:** Department of Basic Science, Graduate School of Arts and Sciences, the University of Tokyo, Meguro, Tokyo; Universal Biology Institute, the University of Tokyo, Bunkyo, Tokyo

**Keywords:** heterosis, evolution, phenotypic variance, gene regulatory network, genetics, sexual reproduction

## Abstract

Heterosis describes the phenomenon whereby a hybrid population has higher fitness than an inbred population, and has previously been explained by either Mendelian dominance or overdominance, where it is generally assumed that one gene controls one trait. However, recent studies have demonstrated that genes interact through a complex gene regulatory network (GRN). Furthermore, phenotypic variance due to noise is reportedly lower for heterozygotes, whereas the origin of such variance-related heterosis remains elusive. Therefore, a theoretical analysis linking heterosis to GRN evolution and stochastic gene expression dynamics is required. Here, we investigate heterosis related to fitness and phenotypic variance in a system with interacting genes, by numerically evolving diploid GRNs. According to the results, the heterozygote population exhibited higher fitness than the homozygote population, that is, fitness-related heterosis resulting from evolution. In addition, the heterozygote population expressed lower noise-related phenotypic variance in expression levels than the homozygous population, implying that the heterozygote population is more robust to noise. Furthermore, the distribution of the ratio of heterozygote phenotypic variance to homozygote phenotypic variance exhibited quantitative agreement with previous experimental results. By applying dominance and over-dominance to the gene expression pattern rather than only a single gene expression, we confirmed the correlation between heterosis and overdominance. We explain our results by proposing that the convex high-fitness region is evolutionarily shaped in the genetic space to gain noise robustness under genetic mixing through sexual reproduction.

**Significance Statement:** Heterosis, that is higher fitness in hybrid populations than inbred populations, is a long-standing problem in genetics, evolution, and breeding studies. Heterosis is not necessarily the result of a trait uniquely determined by a single gene, but likely involves stochasticity and interactions among genes. Through numerical evolution of the gene regulatory network, we demonstrate that heterosis is shaped by evolution to achieve robustness against noise and genetic mixing by sexual recombination in gene expression dynamics. That is, a mixed population is more robust to noise than inbred populations, as revealed experimentally by reduced phenotypic variance. The observed link between heterosis and phenotypic robustness, Mendelian dominance, and the convex single-humped fitness landscape represents a novel avenue in evolution and genetics research.

## Introduction

Sexual reproduction generally involves the mixing of genetic information from multiple parents. Most diploid organisms undergo meiosis, which recombines genomes between parents to produce gametes, then combines the gametes to produce the offspring’s genome. One of the most remarkable phenomena in sexual reproduction among populations is heterosis, whereby hybrid populations exhibit higher fitness than inbred populations, which are obtained by repeated sexual reproduction among genetically close individuals. The origin of heterosis has been discussed from a genetics perspective (1–4), in terms of Mendelian dominance.

In Mendelian dominance (5), if two diploid parent genomes have the genes *AA* and *aa*, and show the corresponding traits ‘A’ and ‘a’, then the child has the gene *Aa* and shows trait ‘A’ when trait ‘A’ is dominant and trait ‘a’ is recessive. When the gene *Aa* expresses the trait over ‘A’ or ‘a’, it is called over-dominance. For decades, the relationship between dominance (or overdominance) and heterosis has been recognized as a significant genetics problem; The first is the dominance hypothesis, which explains heterosis by assuming that Mendelian dominance cancels out the phenotypes that reduce fitness in heterozygotes. In contrast, the overdominance hypothesis proposes that heterozygotes can have a novel trait that exhibits higher fitness than both parental homozygotes. However, no solid conclusions have yet been reached(6, 7).

In the standard context of heterosis, the fitness or trait is compared between inbred and hybrid populations. In contrast, Phelan and Austad examined data relating to the traits of over a hundred species, and demonstrated that the variance of traits also shows heterosis, in the sense that the variance of traits for hybrid populations is smaller than that of inbred populations (8). They also discussed the potential relevance of genotype-environment interactions for this variance-related heterosis; however, its theoretical explanation remains elusive.

To discuss the variance-related heterosis, we need to consider the noise or environmental variation, and robustness of traits against such variation(9–13). However, the possible link between noise and heterosis has rarely been discussed (14), despite the fact that stochasticity in phenotypes (gene expression) has recently received substantial attention in the context of homeostasis and cell state selection (15–18).

Moreover, the evolution of gene regulatory networks has been extensively studied (19–23). To date, arguments related to heterosis are based on the simple genotype–phenotype relationship, that is, the case in which one gene determines one phenotype (5–7). However, the existence of a complex genotype–phenotype relationship caused by interactions among genes has garnered substantial attention (16). The importance of gene regulatory networks has also been recognized, in which protein expression depends on the mutual activation or suppression of genes, where a phenotype, for example, expression of a single protein, depends on expression levels of multiple genes(19). Recent transcriptome analysis experiments have revealed that such interactions between genes (24), which constitute a regulatory network, determine the expression or non-expression of genes. Furthermore, regulatory interactions (promoters, transcription factor binding sites, enhancers) change more frequently through the evolutionary course than the respective genes that encode specific proteins(25).

For this purpose, we extend the GRN to a diploid case. Although a few studies have addressed heterosis using GRN models (26, 27), the consequences of GRN dynamics and the robustness of phenotypes to noise and genomic mixing by sexual recombination have not previously been addressed. Therefore, we analyze these topics by simulating the evolution of diploid gene expression dynamics and discuss the mechanism of hybrid fitness and phenotypic variance.

The organization of this paper is as follows. First, we introduce the model of the diploid GRN. Then, we compare the fitness of homozygote and heterozygote populations and reveal the fitness-related heterosis. Next, we compare the phenotypic variances between homozygote and heterozygote populations to demonstrate that the latter takes much smaller values. Additionally, we compare the ratio of heterosis related to phenotypic variance obtained in our simulation to previous measurement data (8). We also reveal the correlation between hybrid fitness and the dominance or overdominance of gene expression patterns. Finally, an intuitive explanation of the results is provided in terms of the fitness landscape.

### Brief description of the model

#### Evolution simulation based on a diploid GRN model

First, we introduce the theoretical GRN model (We basically follow the model in (13).). Each gene *i*(= 1, 2, …, *N*) has an expression level *x*_*i*_(*t*) at time step *t*. In the model, *x*_*i*_(*t*) is scaled such that it takes a value between zero (non-expression) and one (expression). Each gene interacts with others and itself, with interaction between the *j*th gene and the *i*th gene described by the matrix *J*_*ij*_ (19–23). *J*_*ij*_ can take three values: +1, *−* 1, or 0, which represent the activation, inhibition, and lack of interaction between gene *i* and gene *j*, respectively.

We adopt a discrete-time model (17, 19, 28, 29), in which the expression level *x*_*i*_(*t*+1) in the next time step is determined by 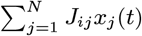, which represents the GRN. Gaussian noise, *η*(0, *σ*) of small magnitude is added to the dynamics to account for the stochasticity in gene expression. The noise term is multiplied by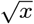 to avoid a negative value of *x*_*i*_(*t*). According to the sigmoid function *f* [*x*] = 1*/*(1 + exp[*− βx*]) with a large *β*(= 100), the expression dynamics are given by

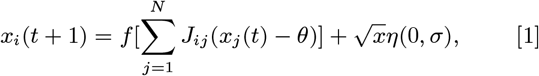

where *θ* is the threshold for the expression level. Starting from the given fixed initial expression levels {*x*_*i*_(0)}, where *x*_*i*_(0) = 0 for most genes and one for several (10) genes. *x*_*i*_(*t*) reaches a fixed point after a certain time *T* in most cases, where *x*_*i*_ is zero or one depending on the structure of the network *J*_*ij*_.

Diploid cells have two genomes that provide two matrices, 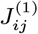 and 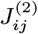. The gene expression dynamics of a diploid GRN are determined by the sum of these matrices; thus, the dynamics are given by modifying 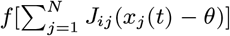 to 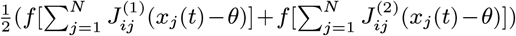 as follows:

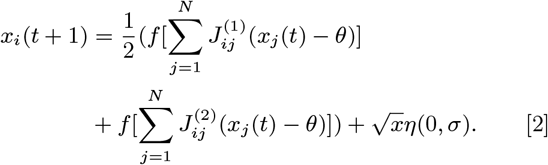

Genetic changes progress according to the following procedure. First, fitness is determined by the distance of the expression pattern of genes {*x*_*i*_(*T*)} from a prescribed target pattern for *i* = 1, 2, …, *M* (*< N*). Two parents are chosen with a probability proportional to the fitness. Two genomes from each parent, 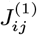 and 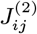, were mixed by recombination to provide a gamete. Next, two gametes from the two parents provide a new 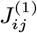 and 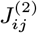 in the next generation. After meiosis, the elements of the new 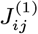 and 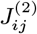 are modified by a mutation with a mutation rate of *μ*. This mutation is included as a change in the matrix element *J*_*ij*_ of 0 or *±* 1 (see Materials and Methods for details on the selection procedure).

#### Defining homozygote and heterozygote populations

The homozygote population is generated as a diploid population consisting of precisely the same diploid genome: 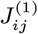 and 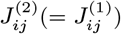. Then, the population of heterozygotes (heterozygote population) is generated as a diploid consisting of two genomes (matrices): 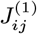 and another 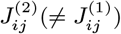.

We first measured the fitness of the homozygote and heterozygote populations, where the mean values for the populations are given by *W* ^homo^ and *W* ^hetero^, respectively. Next, phenotypic variance according to noise *V*_noise_ is defined as the magnitude of the variance of *x*_*i*_(*T*) for individuals with the same genotype but with different noise added to the gene expression dynamics (see Methods for the definition). *V*_noise_ is computed as the average of the variance over genes *i* for homozygote and heterozygote populations, giving 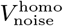 and 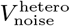, respectively.

## Results

### Fitness-related heterosis

First, we discuss heterosis related to fitness. The change in the degree of fitness-related heterosis *W* ^hetero^*/W* ^homo^ during evolution is shown in Fig.1*A*. The ratio of *W* ^hetero^*/W* ^homo^ increased for earlier generations, and then decreased. After evolution (at the 999th generation), inequality *W* ^hetero^*/W* ^homo^ *>* 1 was maintained; therefore, fitness-related heterosis was generally observed. (See Fig. S1 for the evolution of the distributions of *W* ^homo^ and *W* ^hetero^.) In Fig.1*B*, we plotted *W* ^hetero^ versus *W* ^homo^ for each sample after evolution. Except for three of the 100 samples, *W* ^hetero^ *> W* ^homo^ was satisfied, indicating evolution of heterosis. Then, across each sample, we computed the average ratio of *W* ^hetero^*/W* ^homo^ and examined its dependence on the mutation rate. As shown in Fig.1*C, W* ^hetero^*/W* ^homo^ increased with *μ*, indicating that a greater degree of heterosis evolved at a higher mutation rate. Additionally, at higher *μ*, the ratio of heterosis *W* ^hetero^*/W* ^homo^ was larger at a larger *σ*.

**Fig. 1.**
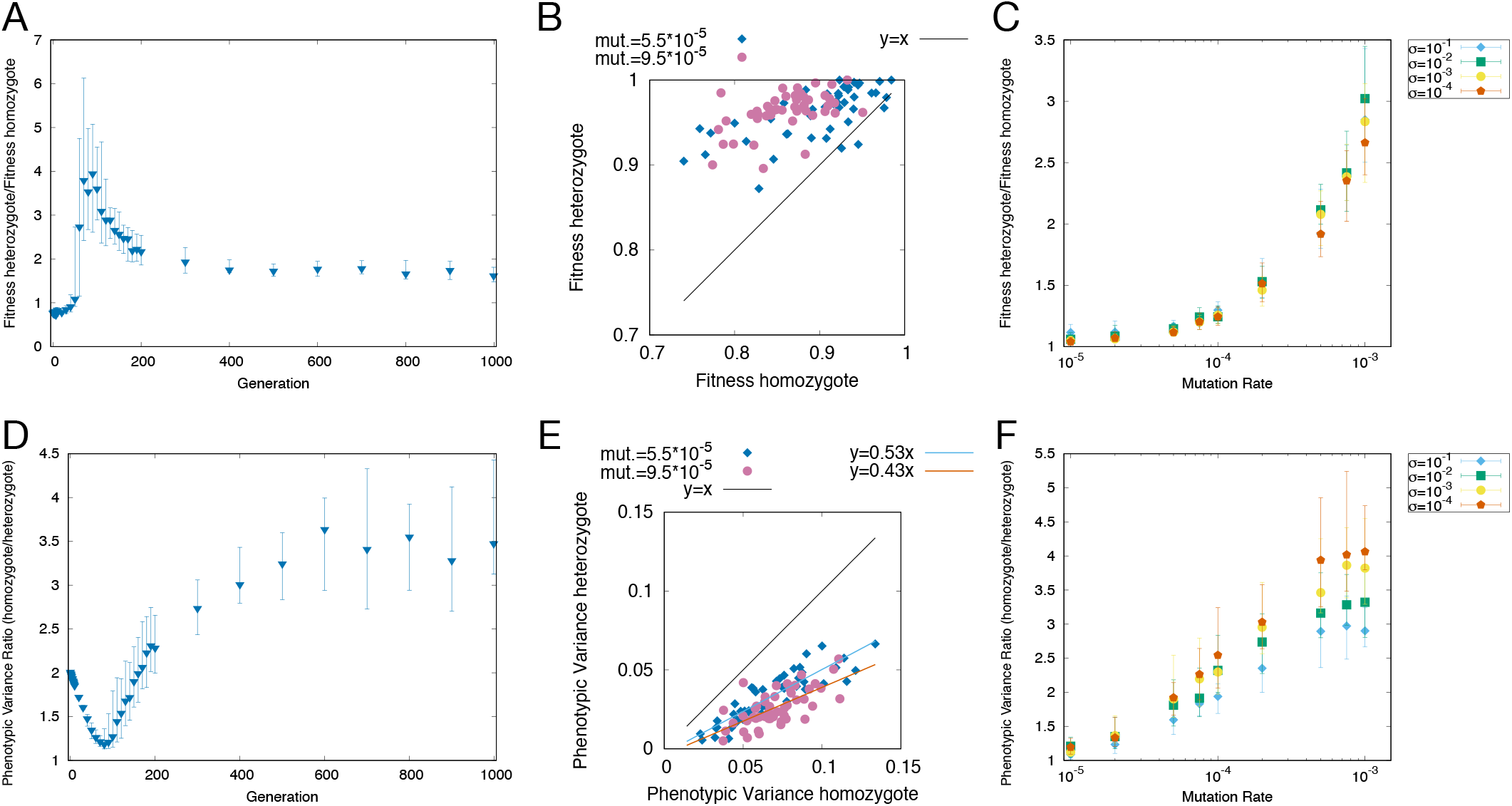
Evolution, variation, and mutation-rate dependence of heterosis. (*A*)Change in the degree of fitness-related heterosis *W* ^hetero^*/W* ^homo^ throughout evolution, calculated for 30 samples in each generation. Points represent the median, and error bars represent the range between the first and third quantiles. *W* ^hetero^*/W* ^homo^ was 0.8 in the 0th generation; it increased initially, then decreased thereafter. The mutation rate per edge was *μ* = 3.0 *×* 10^−4^ and the noise strength was *σ* = 1.0 *×* 10^−4^. (*B*)Variation between *W* ^hetero^ and *W* ^homo^ for each sample. The noise was set to *σ* = 1.0 *×* 10^−4^. Even under the same conditions, variation occurred in *W* ^hetero^ and *W* ^homo^. For all samples in the *μ* = 9.5 *×* 10^−5^ condition, and all except three samples out of 50 in the *μ* = 5.5 *×* 10^−5^ condition, the relationship *W* ^hetero^ *> W* ^homo^ was maintained despite sample-to-sample variation. (*C*)Increase in *W* ^hetero^*/W* ^homo^ with mutation rate *μ*. Points represent the median, and error bars represent the range between the first and third quantile for 30 samples for different noise magnitudes *σ*. The ratio *W* ^hetero^*/W* ^homo^ increased with *μ* and *W* ^hetero^*/W* ^homo^ *>* 1 was maintained. (*D*)Evolution of the mean value of the degree of variance-related heterosis 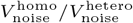. The 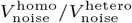 increased with evolution, reaching a median value of approximately 3.0 (*μ* = 3.0 *×* 10^−4^ and *σ* = 1.0 *×* 10^−4^). (*E*) 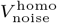 and 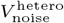 variation for each sample for *μ* = 5.5 *×* 10^−5^ and *μ* = 9.5 *×* 10^−5^. The relationship 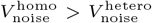 was maintained despite sample-to-sample variation. (*F*)Change in 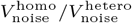 with *μ*. 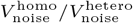 increased with *μ* in this range.

### Variance-related heterosis

The evolution of the mean value of the degree of variance-related heterosis 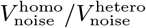 is shown in Fig.1*D* (see Fig. S2 for the evolution of the distributions of 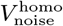 and 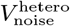). 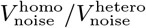 increased from two at the 0th generation to approximately three after evolution (at the 999th generation). Note that, for a random network, the estimated ratio is two (see SI Appendix). Hence, the results indicate that evolution created non-trivial heterosis related to variance. We call this phenomenon variance-related heterosis to distinguish it from fitness-related heterosis. Because a lower variance *V*_noise_ corresponds to higher robustness to noise, the phenotype of heterozygous diploids is more robust against noise than homozygotes.

As shown in Fig.1*E*, we measured 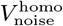 and 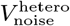 after evolution for different samples under a given mutation rate and noise magnitude. In all samples, 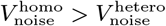, which indicates that fluctuation of the heterozygote was smaller than that of the heterozygote. The dependence of 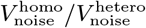 on *μ* is shown in Fig.1*F*. We confirmed that 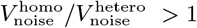 for all *μ*, whereas the ratio increased with *μ*. For *μ >* 5 *×* 10^−5^, 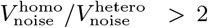 was obtained after evolution, showing higher variance-related heterosis than for the case of random networks. At higher *μ*, the ratio of heterosis 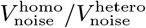 was larger at a larger *σ*.

### Comparison with previous measurement data

By measuring the variance of traits from data of 172 traits drawn from 15 different species, Phelan and Austad reported that hybrid populations exhibited fewer phenotypic fluctuations than inbred populations (8). These data are explained in (8) as follows: *“Data covering a broad range of taxa, including invertebrate and vertebrate animals as well as plants, demonstrate that the assumption that genetic variability and environmental variability are independent of each other is unwarranted*.*”* The values listed are the phenotypic variance of the inbred population and crossbred population divided by the mean (coefficient of variation, CV). Note that the variance adopted in (8) is not equal to the phenotypic variance by noise (*V*_noise_). The variances adopted in their data also include that due to the genetic variation. However, as the *V*_noise_ and the all phenotypic variance are positively correlated (proportional) in previous studies (30–34). Hence, we compare the results of *V*_noise_ with the variance data in (8).

We compared the ratio of phenotypic variances for inbred and crossbred populations in the previous experimental data by the 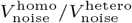 value obtained numerically in our model. We then plotted the distribution of 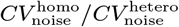 in Fig.2, and 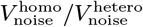 for a low mutation rate in our simulation. The two distributions exhibit similar quantitative values, implying that the variance-related heterosis obtained by the diploid GRN model is consistent with the field data.

**Fig. 2.**
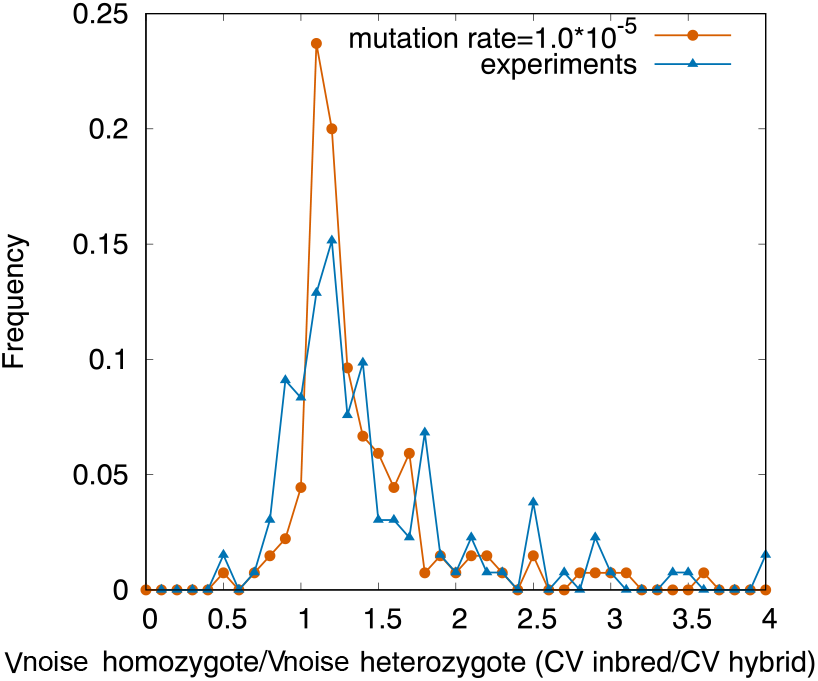
Distribution of the ratio of variance (CV) of 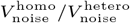 for our model and previous experimental data. The simulation results (orange dots) are for *μ* = 1.0 *×* 10^−5^, whereas the measurement data for *CV* ^inbred^*/CV* ^hybrid^ are measured for inbred and crossbred populations (blue triangles) from (8).

### Correlation of heterosis with pattern dominance and pattern overdominance

In this subsection, we discuss how heterosis is correlated with dominance and overdominance, as defined in (6, 7). Here, we need to extend the definition of dominance and overdominance to adapt to the case in which a collective of expressed genes in a network determines the phenotypic traits, rather than a single gene. Thus, we define and compute pattern dominance and pattern overdominance from the gene expression patterns and explore their correlation with heterosis(13).

To introduce the dominance of gene expression patterns, we first compute the gene expression patterns over a group of genes for given homozygotes *AA* and *BB* taken from a given genome pool, which is represented by 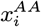 and 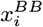. Next, we compute the gene expression pattern of heterozygotes *AB*, given by 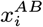, for selecting the genes 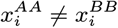. Then, we examine whether the expression pattern of the heterozygote is biased toward one of the homozygote expression patterns, for example, 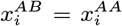 or 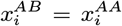, and compute the fractions. Pattern dominance is defined by a larger between the two. If the expression of each gene is independent, such a bias is not expected, and the pattern dominance is 0.5. The pattern dominance is one if 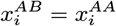 or 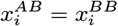 for all *i* with 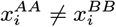. In contrast, for a set of genes with 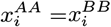, the fraction of such genes *i* of heterozygotes satisfying 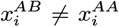 gives the degree of pattern overdominance (see the Methods section for the definitions of pattern dominance and pattern overdominance).

The correlation between pattern dominance, pattern over-dominance, and fitness-related heterosis is shown in Fig.3 (see Fig.S3 for the correlation with variance-related heterosis). Heterosis *W* ^hetero^*/W* ^homo^ exhibited a negative correlation with pattern dominance but a positive correlation with pattern over-dominance. The positive correlation between fitness-related heterosis and pattern overdominance seems natural because the expression of heterozygotes must be different from that of homozygotes in order to achieve heterosis. On the other hand, heterosis was previously explained by dominance, because dominance can compensate for the deleterious mutations in multiple loci. However, the negative correlation between fitness-related heterosis and pattern dominance, observed in our simulation, supports the overdominance explanation rather than the dominance explanation.

**Fig. 3.**
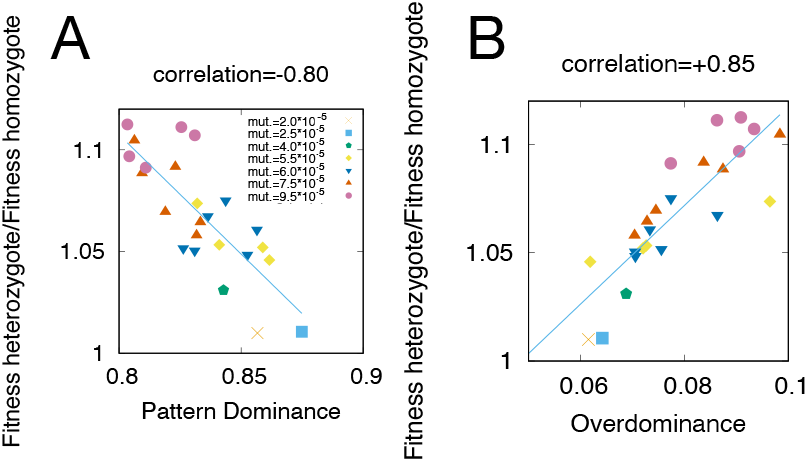
Correlation between heterosis and dominance (overdominance). (*A*) Correlation between pattern dominance and *W* ^hetero^*/W* ^homo^, with a correlation coefficient of −0.80. (*B*) Correlation between pattern overdominance and *W* ^hetero^*/W* ^homo^, with a correlation coefficient of +0.85. Each point corresponds to different mutation rates denoted by different symbols, whereas different points correspond to different noise magnitudes *σ* of [5 *×* 10^−5^, 1 *×* 10^−2^]. Points represent the average of 50 samples, calculated only for the 999th generation out of 50 samples with a fitness average of 0.9 or higher.

Similarly, variance-related heterosis was negatively correlated with pattern dominance but positively correlated with pattern overdominance; however, the correlation coefficient was smaller than that for fitness-related heterosis (see Fig. S3).

### Explanation of the results according to the convex fitness region and single-humped fitness function

In this section, we suggest an intuitive explanation of our results using fitness landscape pictures. Nevertheless, further mathematical elucidation is required in the future.

We consider a fitness landscape over a genotype space (35). In particular, we discuss the high-fitness region. After evolution, offspring are produced from the two parents by meiosis within this high-fitness region. The genotypes of the diploid genome 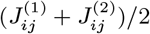 will be located approximately at the internal division of the genotype of the parents. The offspring must have high fitness; otherwise, the offspring will not be selected. For example, when the higher-fitness region is not convex (Fig.4*A*), the offspring may not be selected because of their lower fitness compared to the parents. Therefore, the high-fitness region for the population under sexual reproduction and selection shapes the convex region in the genotype space (Fig.4*B*).

**Fig. 4.**
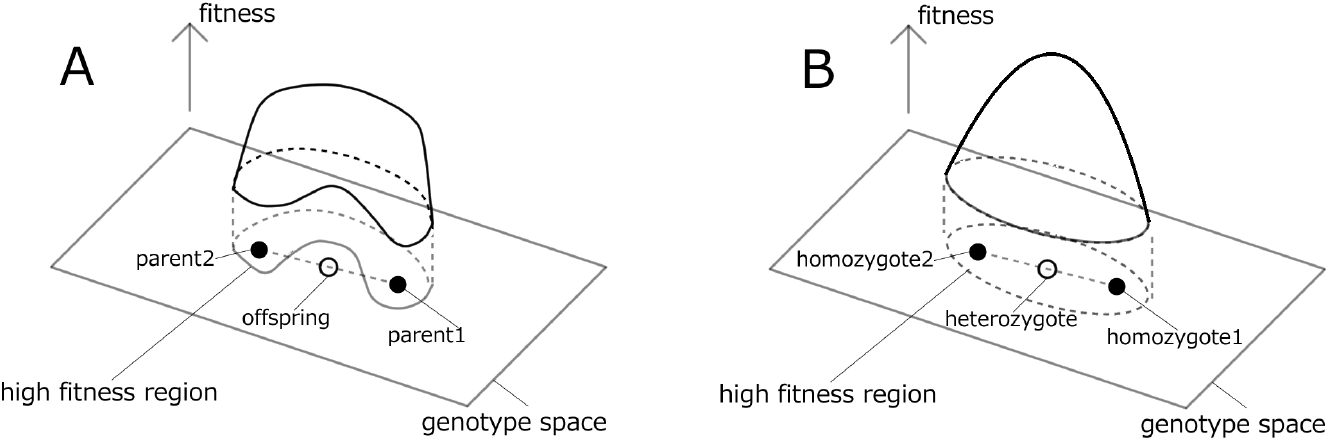
Schematic of the fitness landscape and high-fitness region. (*A*)When the high-fitness region is not convex, the offspring may have lower fitness than the parents; therefore, the linage will not be selected. (*B*)When the high-fitness region is convex over the genotype space, the offspring’s fitness is higher than or equal to that of the parents. Thus, the fitness of the heterozygote is higher than that of the homozygote, leading to heterosis.

The fitness landscape, then, is expected to take a single-humped structure around the center of the region that decreases as the genotype moves toward the periphery of the high-fitness region (Fig.4*B*). When we consider the robustness of the fitted state, the fitness is expected to be less sensitive to genetic change around the center than at the periphery. This suggests that the fitness function is flat around the center and steeper farther from the center (i.e., convex, single-humped functions). Thus, the variance of the fitness (phenotype) will be smaller around the center.

Now, consider the distribution of the homozygote population in a high-fitness region, whose genotype can be broadly located. This population contains individuals with low fitness and large phenotypic variance. Therefore, a heterozygote population generated from a pair from the homozygote population is biased toward the center of the region. Accordingly, the distribution of the heterozygote population is expected to have higher fitness and lower variance than that of the homozygote population, leading to fitness- and variance-related heterosis. This picture is also consistent with the observed correlation between heterosis and overdominance.

Thus, we expected the fitness landscape to be single-humped with a flatter top (i.e., convex), and examined the validity of this expectation in our model. First, we computed the distance between the two genome matrices 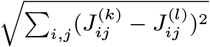 for the two genomes (*k, l*) in the population of the 999th generation. Among the combinations, we chose unfitted genomes, that is, the homozygotes of both *K, L* genomes had fitness *≤* 0.95 and the largest distance between genome matrices 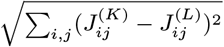. Next, we produced 100 matrices corresponding to the internal division of 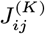 and 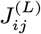, that is, 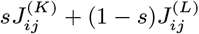, by changing *s* step-by-step in the range 0 *≤ s ≤* 1. Then, we computed the mean fitness for homozygotes over 100 matrices of each division point given by *s*.

As shown in Fig.5, the fitness function exhibited a single-humped structure against the internal division parameter *s* and had a flatter landscape around the top, as predicted by our proposed explanation. As this type of fitness function is not observed in random matrices *J*_*ij*_ (corresponding to the 0th generation), we suggest that it was acquired by evolution during sexual reproduction. This fitness increase at the center is further enhanced by introducing differences in the genomes in diploid (heterozygosity) with a few percentages.

**Fig. 5.**
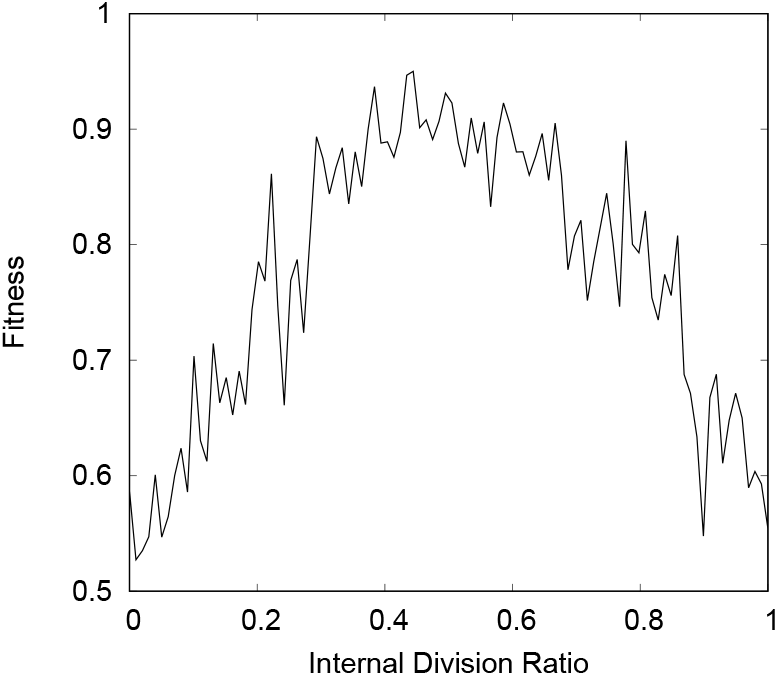
Fitness according to changes along a given genome pair, parametrized by *s*, where the internal division genome matrix is given by 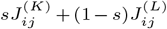 between genome pairs (*K, L*). This was chosen so that the fitness for (*K, L*) was less than 0.95 after evolution (at the 999th generation) and the distance between genome matrices was the maximum value. The horizontal axis represents the internal division parameter *s*. The average fitness over 30 samples is plotted for *μ* = 3.0 *×* 10^−4^ and *σ* = 1.0 *×* 10^−4^.

## Discussion

In the present study, we employed numerical evolution of the diploid GRN to demonstrate heterosis related to fitness and phenotypic variance, as well as its correlation with overdominance, and compared the simulation results with previous measurement data. First, heterozygote populations exhibited higher fitness than homozygote populations, indicating heterosis. Heterosis exhibited a negative correlation with dominance and a positive correlation with overdominance. Moreover, phenotypic variance due to noise was lower in the heterozygote population than in the homozygote population, indicating variance-related heterosis. Furthermore, the numerically obtained distribution of the ratio of homozygote population variance to heterozygote population variance agreed quantitatively with that of previous measurement data (8).

We then proposed a possible explanation for these results according to the convex fitness landscape. Specifically, the observed landscape formation may arise if high fitness is maintained under the conditions of sexual reproduction and noise. Therefore, to increase or maintain fitness during sexual reproduction, it is necessary to create a population of genomes in which the fitness of the offspring is equal to or greater than that of the parents. In addition, the population located inside the high-fitness region will evolve to avoid the periphery of the region in which the phenotype (fitness) is vulnerable to noise. This fitness landscape implies epistasis. However, epistasis alone is not sufficient to support heterosis. A convex high-fitness region with a fitness function that flattens around the top is a stronger hypothesis than epistasis, as it concerns the global shape of the fitness landscape for the entire genome population. The validity of this hypothesis requires further examination in theoretical models with evolving gene expression dynamics, as well as experimental data.

We also studied the relationship between overdominance and heterosis. Instead of a single locus, the dominance and overdominance of expression patterns at multiple loci were defined and measured in this study. Note that the previous explanation for heterosis by overdominance was based on the locus-specific overdominant effect in (36), whereas its molecular mechanism is still unclear(37). In contrast, pattern overdominance, as defined in this study, is concerned with the robustness of a system with interacting genes(11, 12), whereas future experimental confirmation is needed. Furthermore, because the distribution of the underlying genomic population is diverse, the genomic distribution of the homozygote and heterozygote populations should be measured separately to ensure a careful analysis of heterosis.

Finally, we discussed variance-related heterosis, that is, the relationship between phenotypic variance in heterozygotes and homozygotes. Recall that even a random network can produce considerable variance-related heterosis; however, an evolved population shows stronger variance-related heterosis. The result of variance-related heterosis can also be explained by the single-humped fitness function described above, which increases robustness to noise. Such noise robustness has recently received substantial attention in the context of homeostasis, and cell state selection (15–18, 38); thus, heterosis should also be reexamined in the light of noise robustness.

In this study, our numerical results were compared with data previously compiled by Phelan and Austad (8). We uncovered remarkable similarities in the distributions of the ratio of variance-related heterosis, whereas absolute values were not necessarily the same. The simulation results assumed a given fixed species and fitness condition, whereas the measurements were derived from different organisms under different environmental conditions. In order to make a further quantitative comparison, experiments should be conducted for given species under fixed environmental conditions, with ploidy manipulation and measurement of the noise. For example, both are available in yeast(39–41), in which the GRN has already been analyzed. A quantitative analysis of both heterosis and phenotypic noise is also required in future research.

To conclude, we elucidated the evolution of heterosis in a system of interacting genes, by noting the relevance of robustness to noise as well as sexual recombination(10) and meiosis of genomes(42). This evolution of heterosis also implies advantages of sexual reproduction, which goes beyond the picture of Muller’s ratchet(43) that is a removal of deleterious mutants: The overdominance explanation supported in this study suggests that heterozygosity in diploid organisms is advantageous with interactions between the two genomes, by shaping a convex fitness landscape through evolution. As the present evolution does not assume breeding that is often the target of standard heterosis studies, the present study will be applied for the evolution in the field and laboratory experiments as well.

## Materials and Methods

Let us define 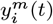 as the concentration of mRNA *i* from each chromosome (*m* = 1, 2) and *x*_*i*_ (*t*) as the concentration of the corresponding protein in a cell. Note that the proteins are synthesized from both chromosomes, where mutations or recombinations are mainly introduced in the promoter or enhancer region, so that the superscript *m* in *x*_*i*_(*t*) is not required. By replicating the regulation in the diploid cell by 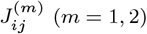 for each gene, we obtain

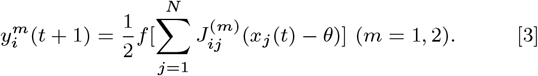

where *θ*(= 0.5) is the threshold of expression, which makes the states *x* = 0 and *x* = 1 symmetric for simplicity. As protein *i* is synthesized from the corresponding mRNA, we then obtain

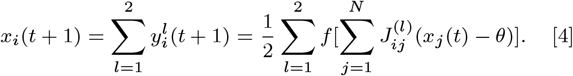

### Evolutional simulation

We take the initial condition *x*_*i*_(0) = 0 for *i* = 1, 2, …, *N*_zero_ and *x*_*i*_(0) = 1 for *i* = *N*_zero_ + 1, …, *N*, where *N*_zero_ = *N −* 10. After a sufficient time, the expression pattern *x*_*i*_(*t*) reaches a stable state *x*_*i*_(*T*). Then, the fitness is determined by the distance between the gene expression levels of the “target genes, “ *i* = 1, …*M* (*< N*). The target pattern is defined as one for all genes, and fixed throughout the evolution. Here, the phenotype is defined as the time average of *x*_*i*_(*t*) for the last 10 steps, 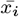. By setting the maximum fitness for 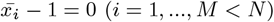, the fitness is defined as 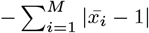. In some cases, the expression pattern *x*_*i*_ (*t*) oscillates over time and cannot reach a stable state; however, these cases are rare. Moreover, after the evolution, such a case is not observed because a stable expression pattern *x*_*i*_ = 1 (*i* = 1, …, *M*) can be more advantageous than an oscillatory state under this fitness landscape. The selection pressure is set to *S*, so that the probability of obtaining the phenotype 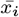 is proportional to 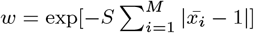.

Two distinct individuals *k*_1_ and *k*_2_ are selected as parents with probabilities of 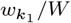 and 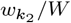, from which the next generation is produced according to the above fitness (*k*_1_ ≠*k*_2_), where *W* is the sum of all fitness values in the population. To introduce meiosis, we adopt the following procedure. A single gamete is generated by mixing 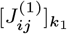 and 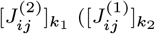and 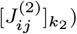 via row vectors with equal probabilities to produce a new genome *n*, where 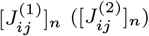.

The simulation is performed by changing the mutation rate (*μ*) is taken in [10^−5^, 10^−2^] and the standard deviation of the noise (*σ*) in [5 *×* 10^−4^, 10^−1^]. The number of genes (*N*) is set to 100, the ratio of non-zero elements in 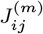, the selection pressure, *S*, is set to 2, and the number of target genes (*M*) is 10. The relaxation time (*T*) to examine the fitness is set to 30. The total number of individuals (*K*) included in the study is 100.

### Definition of *V*_noise_

For a given diploid with 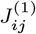 and 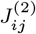, *x*_*i*_ after relaxation is computed under noise conditions. After repeating this process *L* times and obtaining *x*_*i*_ for each run, we measure the variance of the expression pattern *x*_*i*_. The variance *V*_noise_ is computed using the average of the variance over all genes(*N* = 100) and trials (*L* = 100).

### Pattern dominance: definition and calculation

First, we extract 2*N* genomes 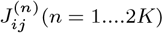 from *K* individuals in the evolutionary simulation. We then make a complete copy for each 2*K* genome, creating a population of 2*K* complete homozygotes, e.g., 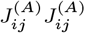. Then, we compute the expression dynamics in each of the 2*K* complete homozygotes to obtain the expression pattern 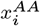. Next, from the 2*K* genomes, two 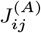 and 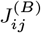 are randomly selected to create a diploid cell consisting of two heterozygous genomes. Then, from the expression dynamics of the heterozygous genomes, the expression pattern of heterozygous 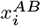 is obtained.

We now compare the expression of genes that are differentially expressed between two such homozygotes, that is, those with 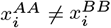. The number of *i* genes with 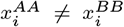 is defined as 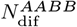. Next, the number of 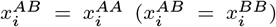 *is* defined as 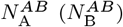. Then, the degree of dominance is given by 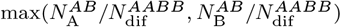.

Note that, in the current model, the expression level of each gene *x*_*i*_ is almost always zero or one. Therefore, if only one gene is differentially expressed between the two homozygotes, the calculated dominance will always be one. To remove this bias, we calculate the pattern dominance only for homozygote 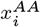 and 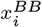 pairs in which more than 30 genes are differentially expressed.

### Pattern overdominance: definition and calculation

Whereas pattern dominance focuses on the expression patterns of genes that differ between homozygotes of the two genomes, pattern overdominance focuses on genes expressed in the same manner between the two genomes. Let 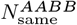 be the number of genes with the same expression as the two pure strains of genomes, and 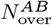 be the number of genes in heterozygote *AB* that are expressed differently from the pure strains *AA* and *BB*. 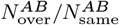 is defined as the degree of pattern overdominance (Fig.6).

**Fig. 6.**
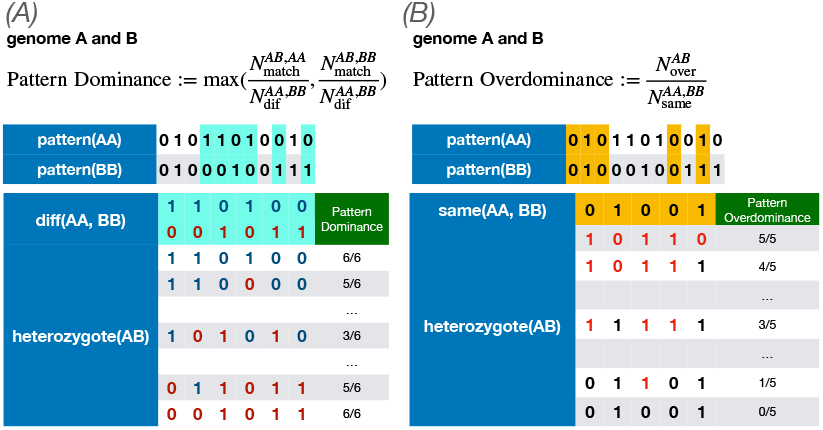
Definition and calculation of pattern dominance and pattern overdominance. (*A*)Pattern dominance: First, we compute the expression patterns of homozygotes *AA* and *BB*. Next, the expression pattern is computed by making the heterozygote *AB*. We then calculate the percentage of the expression pattern obtained from the heterozygote that matches only one of *AA* or *BB* for the differentially expressed genes among the two homozygotes, focusing on the differentially expressed genes in the two homozygotes (blue). Then, the pattern dominance is computed as a larger percentage of matches between 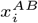 and 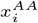 or between 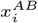 and 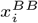. (B) Pattern overdominance: For the gene that shows the same expression in the two homozygotes, we calculate the percentage of the expression pattern obtained from the heterozygote that is different from that of *AA* and *BB*, focusing on the genes that show the same expression in the two homozygotes (blue). Then, the pattern overdominance is computed as the percentage of 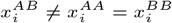.

## ACKNOWLEDGMENTS

The authors would like to thank Tet-suhiro Hatakeyama, Yuichi Wakamoto, Naoki Irie, Akira Sasaki, and Shuji Ishihara for their stimulating discussion, and Daniel Hartl for informing us of (8). This research was partially supported by a Grant-in-Aid for Scientific Research (A) 431 (20H00123) and a Grant-in-Aid for Scientific Research on Innovative Areas (17H06386) from the Ministry of Education, Culture, Sports, Science and Technology (MEXT) of Japan.

