## Supplemental Information for "Heterosis of Fitness and Phenotypic Variance in the Evolution of Diploid Gene Regulatory Network"

**This PDF file includes:**

Supplementary text
Figs. S1 to S3 (not allowed for Brief Reports)

### Supporting Information Text

#### Estimation of $V_{\mathrm{noise}}^{\mathrm{homo}} \approx 0.250$ and $V_{\mathrm{noise}}^{\mathrm{hetero}} \approx 0.125$ in random networks

Here, we explain $V_{\mathrm{noise}}^{\mathrm{homo}} \approx 0.250$ and $V_{\mathrm{noise}}^{\mathrm{hetero}} \approx 0.125$ in random networks, which correspond to the population of the 0th generation in our simulation.

First, recall that the dynamics of $x_i(t)$ are given by

$$x_i(t+1) = f[\sum_{j=1}^{N} J_{ij}^{(1)} x_j(t)] + f[\sum_{j=1}^{N} J_{ij}^{(2)} x_j(t)] + \sqrt{x}\eta(0,\sigma). \quad [1]$$

We assume that, in a random network, the final state (fixed point) is randomly determined. In this case, because homozygotes have the same $J_{ij}$, they can only take two states, $(f[\sum_{j=1}^{N} J_{ij}^{(1)} x_j(t)], f[\sum_{j=1}^{N} J_{ij}^{(2)} x_j(t)]) = (0,0),(1,1)$. Therefore, the normalized expression level is randomly zero or one. The frequencies of the expression levels 0, 0.5, and 1 are 0.5, 0, and 0.5, respectively. On the other hand, in heterozygotes, the expression does not have to be identical in the two genomes; thus, there are four possible states: $(f[\sum_{j=1}^{N} J_{ij}^{(1)} x_j(t)], f[\sum_{j=1}^{N} J_{ij}^{(2)} x_j(t)]) = (0,0),(0,1),(1,0),(1,1)$. The frequencies of the expression levels 0, 0.5, and 1 are 0.25, 0.5, and 0.25, respectively. Therefore, we obtain $V_{\mathrm{noise}}^{\mathrm{homo}} = 0.250$ and $V_{\mathrm{noise}}^{\mathrm{hetero}} = 0.125$.

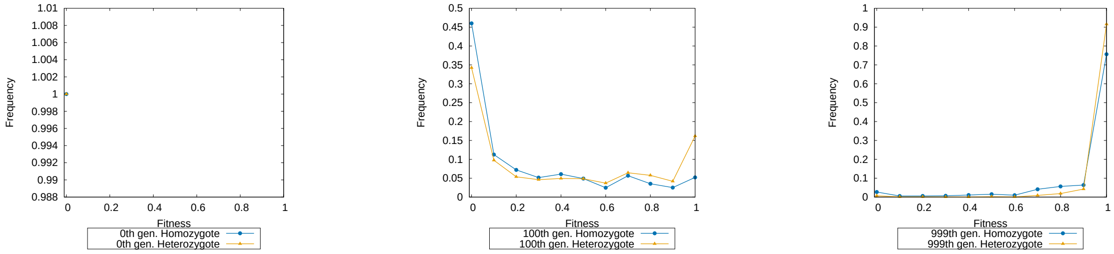


**Fig. S1.** Distributions of $W^{\mathrm{hetero}}$ and $W^{\mathrm{homo}}$ at evolutionary generation 0, 100, and 999. Mutation rate per edge is $\mu = 3 \times 10^{-5}$ and noise strength is $\sigma = 5 \times 10^{-4}$. Bin is 0.1. The distribution is computed for 50 samples. At the 0th generation, a random network cannot achieve expression level one in the target gene; both $W^{\mathrm{homo}}$ and $W^{\mathrm{hetero}}$ are concentrated at zero. At the 100th generation, some of $W^{\mathrm{hetero}}$ exhibit larger values than $W^{\mathrm{hetero}}$. At the 999th generation (after evolution), $W^{\mathrm{hetero}}$ reaches the maximum fitness of close to one more frequently than $W^{\mathrm{homo}}$. Here, $< W^{\mathrm{homo}} ><< W^{\mathrm{hetero}} >$.

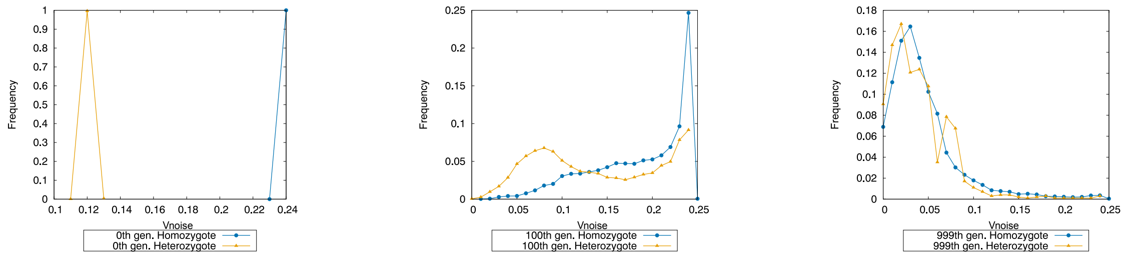


**Fig. S2.** Distributions of $V_{\mathrm{noise}}^{\mathrm{homo}}$ and $V_{\mathrm{noise}}^{\mathrm{hetero}}$. Mutation rate per edge is $\mu = 3 \times 10^{-5}$ and noise strength is $\sigma = 5 \times 10^{-4}$. Bin is 0.01. The distribution is computed for 50 samples. At the 0th generation, the distribution is concentrated at $V_{\mathrm{noise}}^{\mathrm{homo}} \approx 0.250$ and $V_{\mathrm{noise}}^{\mathrm{hetero}} \approx 0.125$, as explained by the random networks. At the 100th generation, the distribution of $V_{\mathrm{noise}}^{\mathrm{hetero}}$ has a much smaller peak value than $V_{\mathrm{noise}}^{\mathrm{homo}}$. Throughout evolution, both $V_{\mathrm{noise}}^{\mathrm{homo}}$ and $V_{\mathrm{noise}}^{\mathrm{hetero}}$ decrease, and the peaks of the distributions are shifted to smaller values. The relationship $V_{\mathrm{noise}}^{\mathrm{hetero}} < V_{\mathrm{noise}}^{\mathrm{homo}}$ is maintained. This implies that the increase in robustness is more prominent for heterozygotes, leading to heterosis related to phenotypic variance.

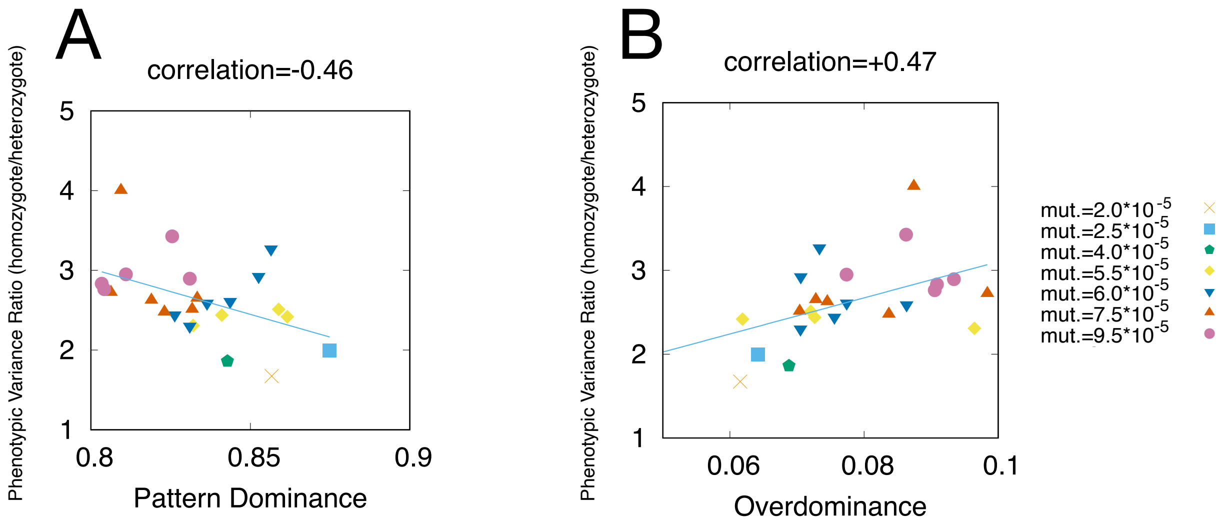


**Fig. S3.** Correlation between heterosis related to phenotypic variance and dominance (overdominance). (*A*) Correlation between pattern dominance and $< V_{\mathrm{noise}}^{\mathrm{homo}}/V_{\mathrm{noise}}^{\mathrm{hetero}} >$, with a correlation coefficient of -0.46. (*B*) Correlation between pattern overdominance and $< V_{\mathrm{noise}}^{\mathrm{homo}}/V_{\mathrm{noise}}^{\mathrm{hetero}} >$, with a correlation coefficient of +0.47. Separate points in each mutation rate series represent different points of noise magnitude $\sigma$ of $[5 \times 10^{-5}, 1 \times 10^{-2}]$. Points represent the average of 50 samples, calculated only for the 999th generation out of 50 samples with a fitness average of 0.9 or higher. A negative correlation is observed between pattern dominance and $< V_{\mathrm{noise}}^{\mathrm{homo}}/V_{\mathrm{noise}}^{\mathrm{hetero}} >$, with a correlation coefficient of -0.46, which is weaker than that between heterosis related to fitness and pattern dominance. Conversely, a positive correlation is observed between pattern overdominance and $V_{\mathrm{noise}}^{\mathrm{homo}}/V_{\mathrm{noise}}^{\mathrm{hetero}}$, with a correlation coefficient of +0.47. Thus, heterosis related to phenotypic variance is achieved as pattern overdominance is acquired; however, the absolute value of the correlation coefficient is smaller than that for heterosis related to fitness .
